# Fibril formation rewires interactome of the Alzheimer protein Tau by π-stacking

**DOI:** 10.1101/522284

**Authors:** Luca Ferrari, Riccardo Stucchi, Aikaterini Konstantoulea, Gerarda van de Kamp, Renate Kos, Willie J.C. Geerts, Friedrich G. Förster, Maarten A.F. Altelaar, Casper C. Hoogenraad, Stefan G.D. Rüdiger

## Abstract

Aggregation of the Tau protein defines progression of neurodegenerative diseases, including Alzheimer’s Disease. Tau assembles into oligomers and fibrils. The molecular basis of their toxicity is poorly understood. Here we show that π-stacking by Arginine side chains rewires the interactome of Tau upon aggregation. Oligomeric nano-aggregates scavenge the COPI complex, fibrils attract proteins involved in microtubule binding, RNA binding and phosphorylation. The aberrant interactors have disordered regions with unusual sequence features. Arginines are crucial to initiate such aberrant interactions. Remarkably, substitution of Arginines by Lysines abolishes scavenging, which indicates a key role for the pi-stacking of the Arginine side chain. The molecular chaperone Hsp90 tames such re-arrangements, which suggests that the natural protein quality control system can suppress aberrant interactions. Together, our data present a molecular mode of action for derailment of protein-protein interaction in neurodegeneration.

**HIGHLIGHTS:**

- Tau fibrils act as fishing net for proteins.
- Tau fibrils attract specific protein families associated with Alzheimer.
- π-stacking by Arginines key for aberrant binding to Tau fibrils
- The Hsp90 chaperone stalls fibril growth and alters interactome

## INTRODUCTION

Protein aggregation is linked to a wide range of neurodegenerative disorders, including Alzheimer’s, Parkinson’s and Huntington’s diseases (Hartl, 2017; Knowles et al., 2014). Protein fibrils are ubiquitous presence in patients’ brains affected by neurodegeneration (Wilcock and Esiri, 1982). For all neurodegenerative disorders, protein aggregation proceeds in a step-wise fashion, from structurally heterogenous oligomeric species to mature fibrils, the former considered to be the most toxic agents (Chiti and Dobson, 2017; Holtzman et al., 2016). Remarkably, it is unclear why protein aggregates are toxic and how they react within the cellular environment of the neuron (Goedert et al., 2017; Wang and Mandelkow, 2016). Intracellular aggregation or the protein Tau is hallmark of Alzheimer’s disease and other fatal tauopathies. (Goedert et al., 2017; Sydow et al., 2011; Wilcock and Esiri, 1982). Several factors play a role in the origin of Alzheimer’s, such as extracellular amyloid formation of the Aβ peptide (Lane et al., 2018). However, intracellular Tau aggregation is sufficient to induce neurodegeneration, correlates with cognitive impairment in humans and is necessary to mediate Aβ toxicty (Bejanin et al., 2017; Roberson et al., 2007; Wang and Mandelkow, 2016).

The mechanistic contribution to disease of Tau oligomers and fibrils remains largely elusive. Due to their structural heterogeneity and difficulties in isolation, it is difficult to point out which cellular processes they target (Holtzman et al., 2016; Wang and Mandelkow, 2016). However, non-physiological protein aggregates may result in new interactions within the cell, disturbing a variety of cellular processes and disrupting the protein quality control network, responsible for fibrils handling and disposal (Ciechanover and Kwon, 2017; Gidalevitz et al., 2006). Major components of this network are two conserved energy-driven chaperone systems, Hsp70 and Hsp90 (Morán Luengo et al., in the press). They both participate in Tau clearance in physiological condition (Blair et al., 2013; Fontaine et al., 2015), however their contribution to neurodegeneration is still elusive. Hsp70 can disaggregate Tau fibrils, while Hsp90 buffers aggregation prone stretches of Tau (Ferrari et al., 2018; Karagöz et al., 2014; Pratt et al., 2015). Hsp90 decreased efficiency during aging may contribute to Tau aggregation, which in turn may dictate further collapse of chaperones activity (Shelton et al., 2017).

Previous proteomics studied revealed that Tau fibrils establish new abnormal interactions with either the insoluble proteome (Donovan et al., 2012) or ER-associated protein complexes (Meier et al., 2015). We recently showed that toxicity of an aggregation-prone variant of the protein Axin is caused by aberrant interactions established by oligomeric nano-aggregates formed in the cytoplasm (Anvarian et al., 2016). Together, this made us wonder how Tau fibrils would interact with the soluble cytoplasmic component of the brain, where protein aggregation takes place. It is an open question whether common structural features govern the binding of interactors to Tau fibrils. It is of great interest to understand which interactions are engaged by Tau fibrils and their binding mechanisms, as this would reveal which cellular processes are targeted by Tau-dependent neurodegeneration and would offer novel therapeutic strategies.

We set out to understand at molecular level how aggregation of Tau modulates interactions with proteins. Here we reveal that fibril formation rewires the interactome of Tau. Interestingly, the avidity properties gained upon fibril formation attract a defined, new set of interactors, which contains disordered regions with a unique amino acidic footprint. The key determinant for derailing the protein network is Arginine-driven π-stacking. This provides a molecular framework to understand aberrant interactions of neurotoxic fibrils. Interestingly, there is a cellular defence system suppressing Tau aggregation. We find that the Hsp90 chaperone stalls formation of Tau fibrils and reshapes their abnormal interactome. Thus, neurons are endowed with a powerful molecular machine that is able to counteract formation of Tau fibrils and their engagement with aberrant interactors.

## RESULTS

### Separation of Tau monomers, oligomers and fibrils

To analyse potential interactome changes upon aggregation of Tau, it is crucial to precisely control fibril formation over time. We set out to biochemically characterise the aggregation process, from monomeric Tau at the start to mature fibrils as end-point. To this mean, we recombinantly produced a pro-aggregating variant of Tau-RD (Tau-Q244-E372, ΔK280 (Gustke et al., 1994)), with a FLAG tag to facilitate detection (Tau-RD*). Tau-RD is an established Alzheimer model that aggregates more aggressively that the wildtype full length protein (Mocanu et al., 2008). We induced its aggregation via heparin following an established procedure (Goedert et al., 1996) and collected samples as aggregation proceeded. We then resolved these samples on density gradients spread over 12 fractions and detected them with a fluorescent antibody **(Fig. 1A).** At time 0, Tau-RD* sedimented on top of the gradient (Fractions 1 and 2, tube I), consistent with its expected monomeric nature at the start of the experiment. After 1 h (tube II), Tau-RD* sedimented further down until fraction 3, indicating the appearance of oligomeric, nanometer-scale aggregates (nano-aggregates). At 8 h (tube III), Tau-RD* spreaded until fractions 7, consistent with the growth of increasingly large aggregates. Finally, after 24 h (tube IV), we detected no Tau-RD* in fraction 1 anymore, indicating that all Tau-RD* aggregated into either oligomers or fibrils, reflected by the presence of aggregates from Electron Microscopy (TEM) characterised Tau-RD* fibrils at 24 h as negatively stained paired helical filaments with periodicity of 50-100 nm, consistent with previous observations (Wegmann et al., 2010) (Fig. 1B). Thus, our procedure generates the full spectrum of Tau species for aggregation-dependent interactome analysis, representing monomers, oligomeric nano-aggregates and mature fibrils.

**Figure 1.**
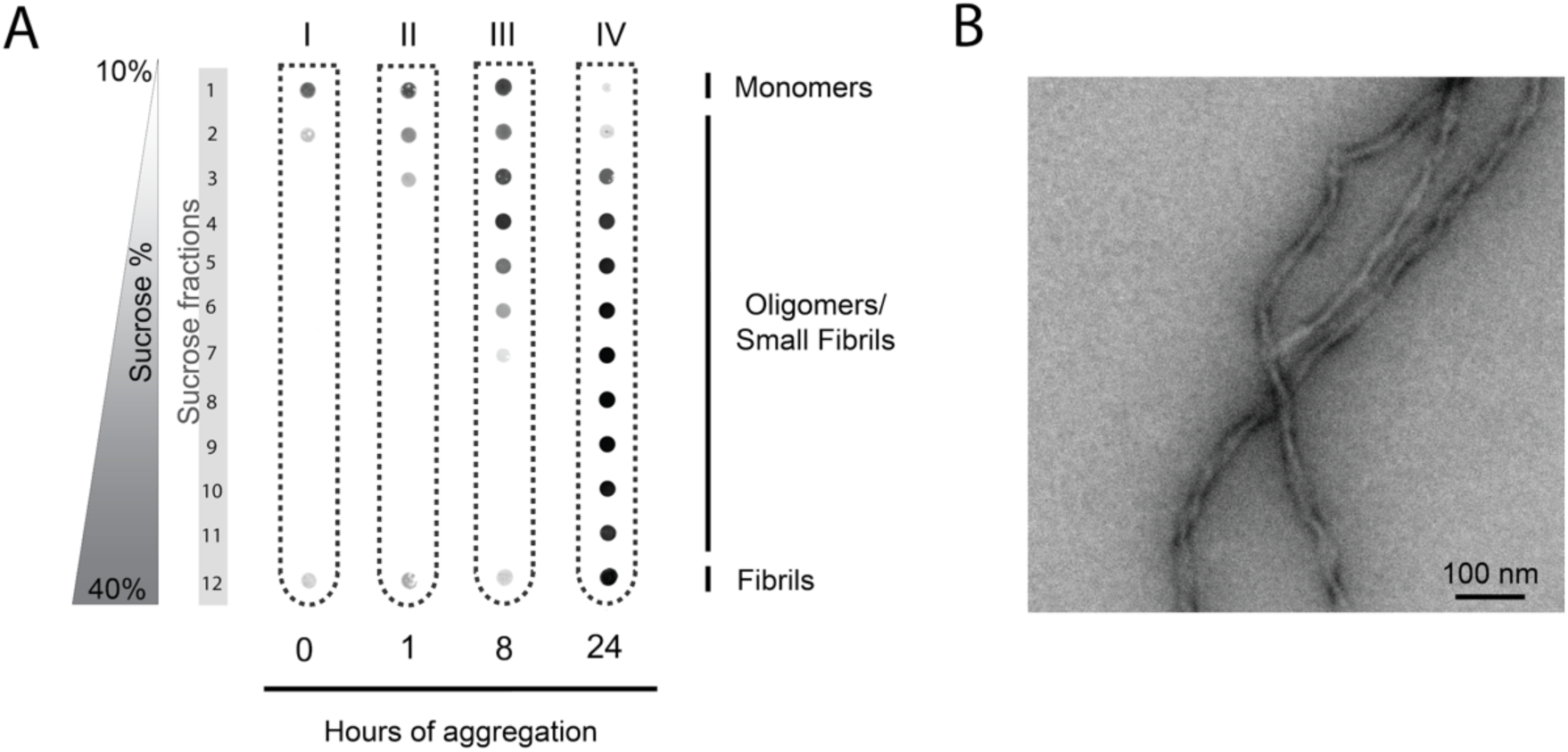
Precise control of Tau aggregation. *(A) Time course of Tau-RD* aggregation monitored by density gradients. Samples were centrifuged for 2.5 h at 200000g.* *(B) Transmission Electron Microscopy of Tau-RD* fibrils, timepoint 24 h. Scalebar 100 nm.*

### Tau fibrils target specific protein families

Next, we wondered how Tau reacts at different aggregation stages with the soluble neuronal proteome. We exposed the FLAG-tagged Tau-RD* at different stages of aggregation during a 24 h period with rat brain lysates depleted of their insoluble components (**Fig. 2A**). We pulled down potential interactors with an anti-FLAG antibody and revealed their identity via mass spectrometry. For each protein, we determined the abundance by counting Peptide Spectrum Matches (PSMs) at each aggregation stage. We compared the protein spectrum at each timepoint (t_x_) to the one of monomeric Tau-RD* without heparin (t_0-_) and plotted changes as logarithmic function. This setup allowed us to monitor proteome changes.

**Figure 2.**
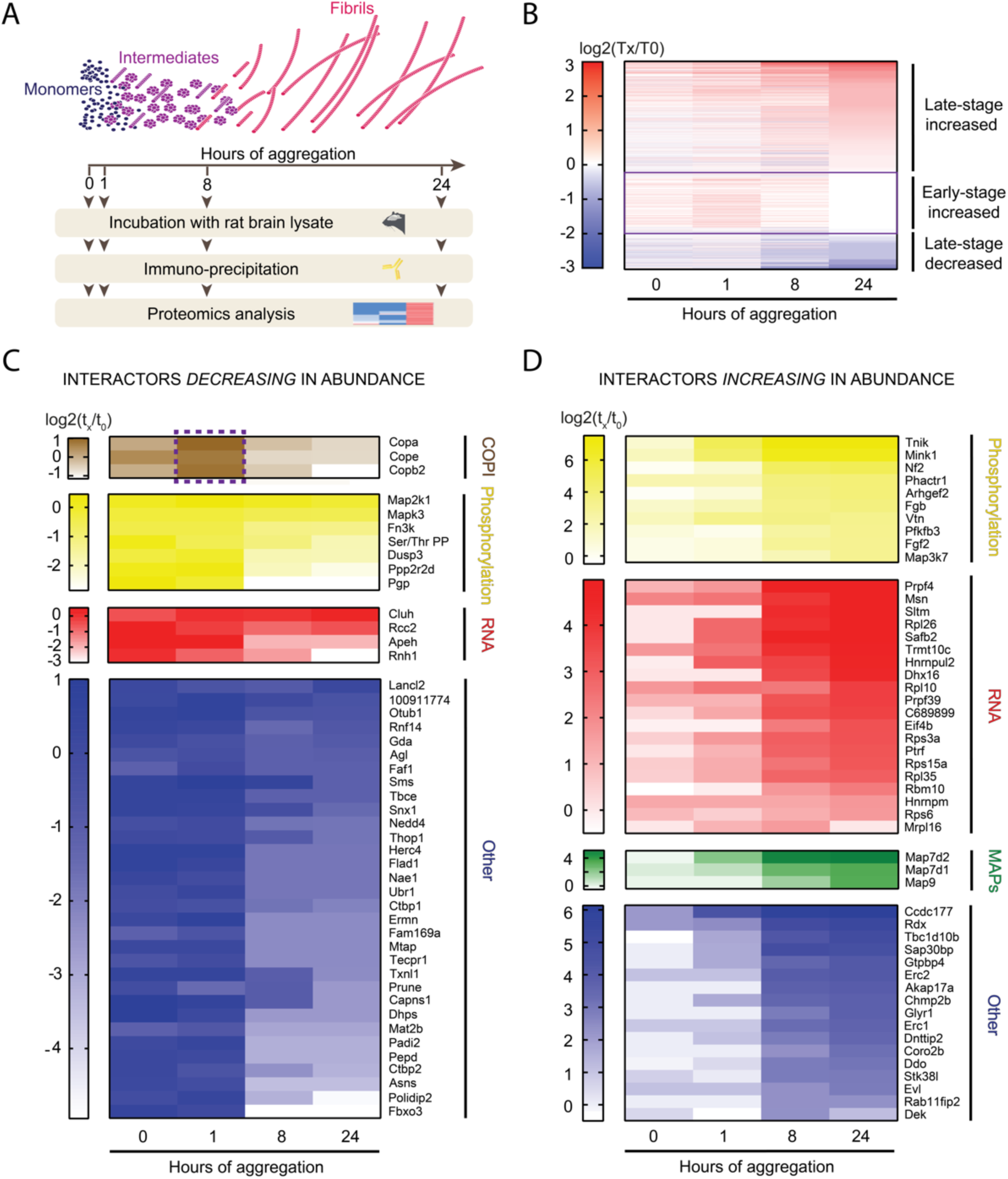
Tau interactome rewires during aggregation. (A) Experimental setup to highlight interactome changes upon Tau aggregation. (B) Heat map of MS-identified proteins of Tau-RD* aggregates (0, 1, 8 and 24 h of aggregation). Colours show relative enrichment of different time points (t_x_) compared to monomeric Tau-RD* without heparin (t_0-_). Interactors are sorted by trend (late-stage decreased, early-stage increased, late-stage decreased). (C) Heat map showing an unbiased selection of Tau-RD* lost proteins (decreasing in abundance) upon aggregation, sorted by functional clusters (GO-term analysis). Dashed purple box highlights interactors increasing in abundance at 1 hour, then decreasing as aggregation proceeds. (D) Heat map showing an unbiased selection of Tau-RD* sequestered proteins (increasing in abundance) upon fibril formation, sorted as for (C).

We uncovered striking interactome changes as Tau-RD* aggregation proceeds (**Fig. 2B**). A sub-group of interactors showed decreased levels compared to the monomeric reference values (**Fig. 2B,** purple box t_24_, blue interactors). Another sub-group of interactors showed binding levels similar for the fibril fractions as for the monomers. Remarkably, however, they bound stronger to early-stage Tau nano-aggregates (**Fig. 2B**, purple box t_1_). Finally, a sub-group of the proteome increasingly associated with Tau upon progression of fibril formation, reaching its maximum as mature fibrils appeared (**Fig. 2B**, purple box t_24_, red interactors). The time-dependent change in protein-protein interactions implies that fibrils and oligomers differ in binding properties to other proteins. Thus, progressive aggregation gradually rewires Tau interactome, with different aggregation species interacting with different molecular partners.

We wondered whether the aberrant interactors of Tau aggregates share common functional properties by classifying them via Gene Onthology enrichment analysis, a tool to cluster proteins with similar biological functions (GO Consortium, 2017 (Ashburner et al., 2000)). We first focused on monomeric-specific interactor lost upon aggregation, with only 25% of interactors belonging to functional clusters, namely protein phosphorylation regulators (16%) or RNA binding proteins (9%) (**Fig. 2C**). Notably, members of the COPI complex are the most prominent interactor of nano-aggregates (t_1_). They neither bind to monomers nor fibrils, indicating that this complex prefers binding to Tau oligomeric species (**Fig. 2C**, dashed purple box). When analyzing the interactome of the Tau-RD* fibrils, remarkably 66% of the newly attracted interactors belong to only 3 major functional clusters **(Fig. 2D),** namely RNA Binding proteins (40% of the interactors), Regulators of Protein Phosphorylation (20%), Microtubule Associated Proteins (MAPs, 6%), with the rest of the proteins showing no GO enrichment (Various Proteins, 34%). Results were confirmed in 3 biological replicates **(Fig. S1** for lost monomeric interactors and **Fig. S2** for fibrillar interactors), allowing us to conclude that Tau aggregates attract proteins connected to a very limited set of biological processes, exchanging proteins connected to the same functional networks and increasing their relative abundance, whereas the rest of the cellular activity seems to not be disturbed by Tau fibrils. Strikingly, COPI preferentially interact with early stage aggregates and MAPs with late stage fibrils, suggesting that different types of aggregates may differ in their gain of function aberrant phenotypes.

### Arginine side chains mediate fibril interactions

We hypothesized that shared biological activity may underline common sequence features. We noted that RNA and microtubule binding proteins and regulatory proteins often contain intrinsically disordered regions. Therefore, we determined the degree of disorder for each fibril-binding protein using the disorder predictor MetaDisorder, which particularly accurate because it is the consensus of 15 primary algorithms (Kozlowski and Bujnicki, 2012). We then plotted the disorder content per functional cluster. All functional clusters, even non-clustered fibrils binders, showed high disorder content, with medians at least 15% higher in absolute disorder than the saverage disordered percentage in the human proteome (Pentony and Jones, 2010) (**Fig. 3A**). Most dramatic is the MAP cluster, which is 95% disordered. We conclude that Tau fibrils attract proteins enriched in intrinsically disordered regions. It is unusual for intrinsically disordered proteins to directly interact with each other (van der Lee et al., 2014), therefore we can also conclude that fibrillation endows Tau with new properties to engage in protein-protein interactions.

**Figure 3.**
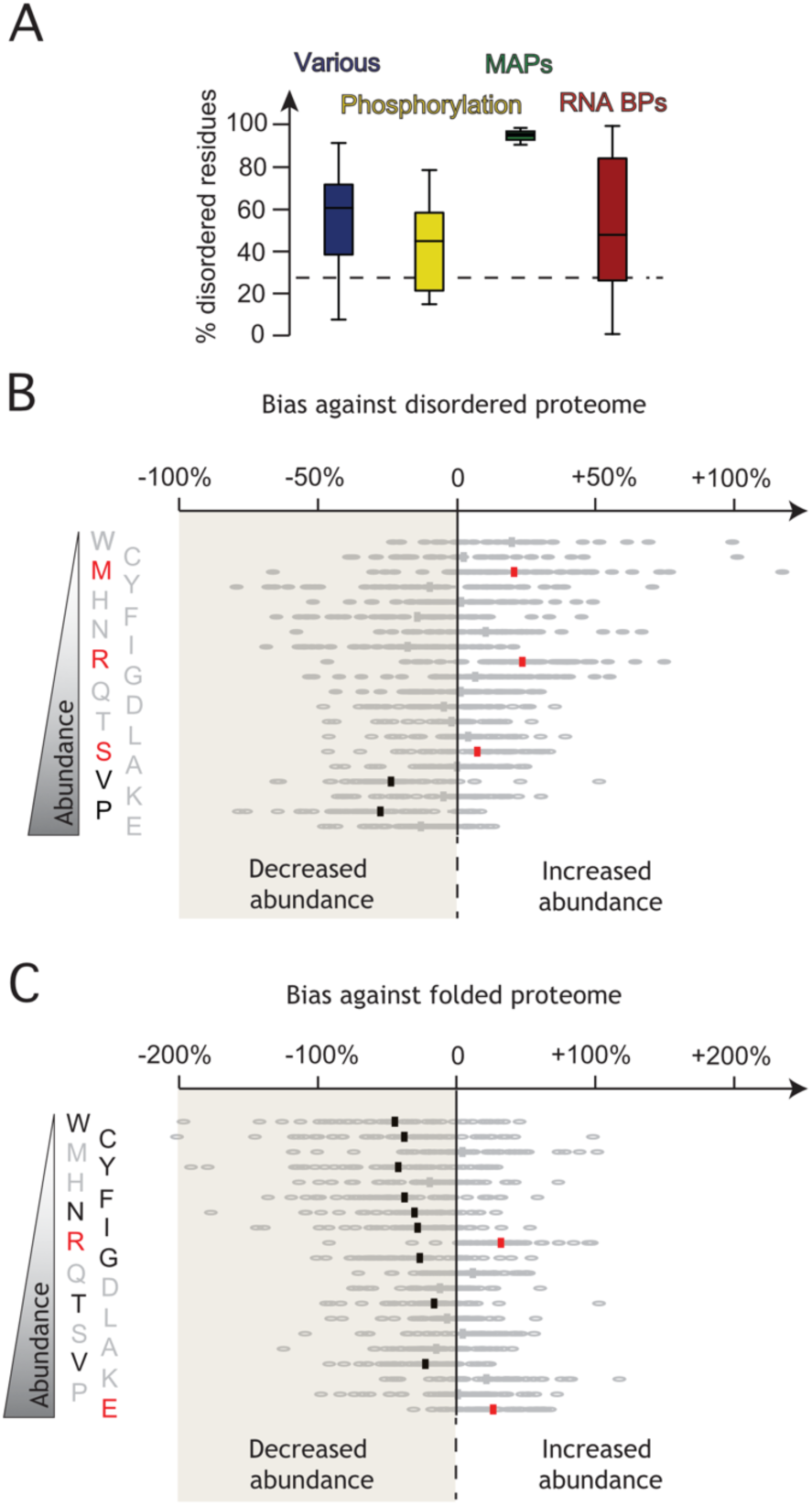
Tau aggregation-specific interactors are enriched in disordered sequences. (A) Prediction of percentage of disordered residues of Tau-RD* fibril-specific interactors (timepoint 24 h) using MetaDisorder (Kozlowski and Bujnicki, 2012), sorted by functional clusters. Dashed line indicates the average disorder percentage for human proteome (Pentony and Jones, 2010). (B) Amino-acidic footprint of disordered regions of Tau-RD* fibril-specific interactors, expressed in fold-changes against the abundance of each amino acid in the disordered regions of the whole human proteome (Tompa, 2002) (red: significantly increased; black: significantly decreased, p ≤ 0.0001). Amino acids on the y-axis are sorted by abundance in the disordered proteome. (C) As in B, but against folded regions of the whole human proteome (Tompa, 2002). Amino acids on the y-axis are sorted by abundance in the disordered proteome, for comparison with Fig. 3B.

We wondered whether disordered stretches captured by Tau fibrils showed specific sequence properties. To reveal such a bias, we compared the frequency of each amino acid in the disordered regions of fibril interactors to the average frequency in the whole disordered proteome (Tompa, 2002). Remarkably, our analysis showed that chemical composition of disordered regions of fibril-specific interactors dramatically differed from that of typical disordered regions (**Fig. 3B**). In particular, Prolines and Valines were significantly depleted, whereas they are the most abundant components in the entire disordered proteome. Conversely, Arginine and Methionine stood out as the most significantly enriched amino acids (p ≤0.0001, Wilcoxon paired non-parmatric t-test). Arginines had higher frequency when compared to Methionines (5% of total residues in disordered regions for Arginine,1.6% for Methionine, **Fig. 3B**). More than 85% of the interactors had at least 1 long disordered region (30+ residues), with 90% of these regions containing at least two Arginines. These numbers reveal that Arginines may have a predominant role in IDRs binding to Tau fibrils. When compared to the folded proteome, most of the amino acids were depleted, (**Fig. 3C**). Conversely, Arginine stood out again as the most significantly enriched amino acid (p ≤ 0.0001, Wilcoxon paired non-parmatric t-test), whereas Methionine had similar enrichment compared to globular proteins, suggesting that the footprint of Tau fibril-specific interactors is different from globular proteins too, and therefore a unique signature of proteins binding to Tau fibrils. Of note, aromatic amino acids, which can also engage in π-interactions but are less well tolerated in disordered stretches, were not enriched when compared with either folded or disordered proteome average frequencies, further highlighting the enrichment of Arginines in binders of Tau fibrils **(Fig. 3B** and **Fig. 3C).** Taken together, our bioinformatics analysis shows that Tau fibril-specific interactors have their own unique footprint, characterized by specific amino acid biases in Arginine-rich disordered regions that differ from the typical disordered segment and globular protein.

### π-π **interactions crucial for fibrils binding**

Next, we aimed to reveal the molecular mechanism of interaction of Tau fibrils with IDRs. Tau-RD has a positive net charge at physiological pH (pI = 9.7 for Tau-RD, 9.3 for Tau-RD*), making us wonder why positively charged Arginines would be so enriched in Tau aggregation-specific interactors. Notably, next to charge-charge interactions, Arginine can also engage in protein-protein interactions via the delocalised π-system of sp^2^-hybridized atoms of its guanidinium group (Vernon et al., 2018). We also noted that there is no significant bias for the other positively charged residue, Lysine (**Fig. 3B**), whose side chains contain only sp^3^-hybridized atoms and are thus unable to engage in π interactions, suggesting that pi orbitals and not charge could be the determinant for binding to fibrils. We set out to test this hypothesis by analysing the impact of Arginine to Lysine exchanges in a fibril-binding protein, an established test to verify the contribution of π-π interactions (Vernon et al., 2018).

We chose the N-terminal domain of the fibril-specific protein Map7 for this test (**Fig. 4**). This domain has extended disordered stretches enriched in Arginines (8.4% of disordered residues), next to a folded coiled coil domain, making it a valuable candidate to test Arginine-driven binding to fibrils. We designed and purified two HA-tagged truncations of Map7-M1-S227, wildtype and an R_12_K variant where we replaced all Arginines in the disordered regions of the protein with Lysines (**Fig. 4A**). The R_12_K substitution had no effect on the net charge of the protein (**Fig. 4B**). Moreover, the circular dichroism spectra of both proteins were identical and they both indicated the presence of α-helixes and disordered stretches (**Fig. 4C**). Thus, exchanging Arginines to Lysines in the disordered segments of the N-terminal fragment of Map7 does neither alter its charge nor its secondary structure.

**Figure 4.**
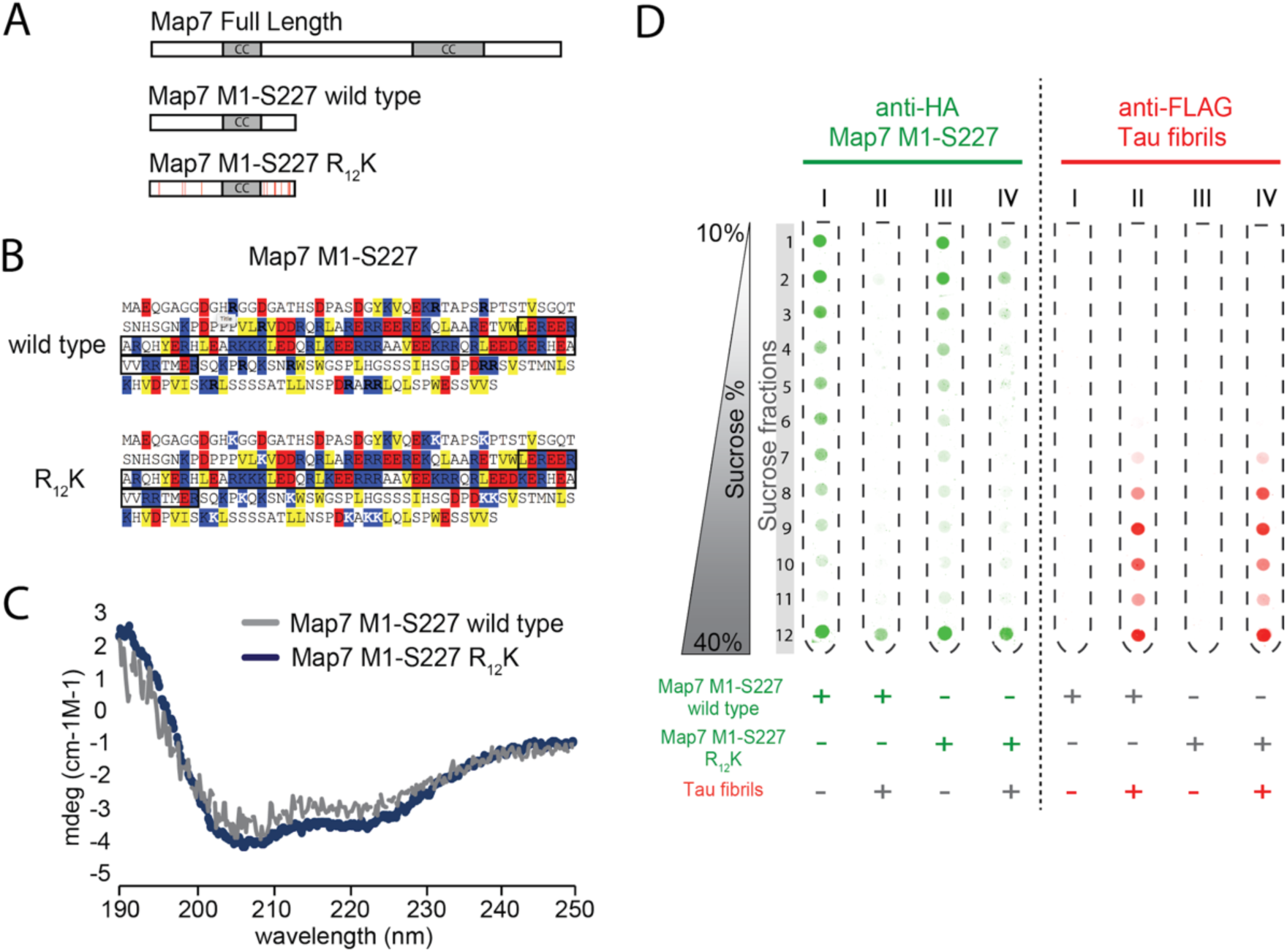
π stacking of Arginines drive binding to Tau fibrils. (A) Graphic scheme of Map7 Full Length and Map7 truncations used in this study. Red bars indicate Rto K replacement. CC = coiled-coil domain. (B) Sequences of Map7 M1-S227 truncations colored by hydrophobicity (yellow boxes), negative charges (red boxes) and positive charges (blue boxes). White Ks indicate Arginines replaced with Lysines. Black boxes comprise coiled-coil domain. (C) Circular Dichroism ofMap7 M1-S227 truncations. (D) Binding of Map7 1M-227S truncations to Tau-RD* fibrils, assessed by density gradients. Tubes (I to IV) were either blotted against HA tag against FLAG tag, targeting respectively Map7 truncations or Tau-RD* fibrils. Samples were centrifuged for 2.5 h at 200000g.

Next, we set out to test whether the exchange of Arginines to Lysines would affect binding to fibrils. We subjected Tau-RD* fibrils to Map7 binding, with or without one of the two Map7 truncations. We resolved protein complexes via density gradients, using different antibodies to detect Tau-RD* and Map7. Map7 wt and R_12_K sedimented throughout the tube (**Fig. 4D**, anti-HA blot, Tube I and III), whereas Tau-RD* fibrils sedimented to the bottom half of the tube (**Fig. 4D**, anti-FLAG blot, tube II and IV, fractions 7-12). The homogenous spread of Map7 alone throughout the tube could be explained by the tendency of disordered regions to show higher apparent molecular mass compered to order structures (Receveur-Brechot et al., 2006). When Map7 wt and Tau-RD* fibrils reacted together, however, the entire Map7 population sedimented in the heaviest fraction, indicating its association with the largest fibrillar structures (**Fig. 4D**, anti-HA blot, tube II, fraction 12), consistent with the proteomics analysis (**Fig. 2C**). Conversely, when Map7 R_12_K and Tau-RD* fibrils reacted, only a part of Map7 population sedimented in the heaviest fraction, whereas another part distributed in the lightest fractions, indicating only partial association with largest fibrillar structures **(Fig. 4D,** anti-HA blot, tube IV, fractions 1-5 and 12). Since both constructs have same charge and structural properties but differ in their ability to engage in π-π interactions, we conclude that π-π interactions are key forces governing binding to fibrils.

### Hsp90 stalls Tau aggregation

Aberrant interactome rewiring is linked to cellular toxicity (Anvarian et al., 2016). It would be therapeutically valuable to find endogenous proteins able to modulate such aberrant transition in interactome. We looked for such endogenous player in the protein quality control network, a pool of proteins that ensures the health of the proteome by removal of misfolded and aggregated proteins (Hartl, 2017). A key player in this network is the molecular chaperone Hsp90, which also controls Tau levels in the cell. Hsp90 forms a complex with Tau buffering its aggregation-prone regions (Karagöz et al., 2014) and controls the degradation of Tau monomers (Dickey et al., 2007).

To understand the effect of Hsp90 on Tau aggregation dynamics, we wondered how this molecular chaperone would interfere with the formation of Tau fibrils. To monitor aggregation of Tau-RD* in the presence of increasing concentrations of Hsp90, we induced aggregation in the presence of Thioflavin T, a dye that recognizes amyloid structures (**Fig. 5A**). After aggregation reached plateau for all conditions, we resolved aggregating samples on density gradients (**Fig. 5B**). In a dose-dependent manner, Hsp90 suppressed Tau-RD* to form higher molecular weight species, decreased amyloid content and prevented fibril formation altogether. Hsp90 also altered morphology of the aggregates: instead of fibrils, only smaller structures were visible in the TEM images when aggregation was performed in the presence of Hsp90 (**Fig. 5C**). Thus, Hsp90 has a dramatic effect on Tau aggregation dynamics, derailing its aggregation toward non-fibrillar nano-aggregates with decreased amyloid content.

**Figure 5.**
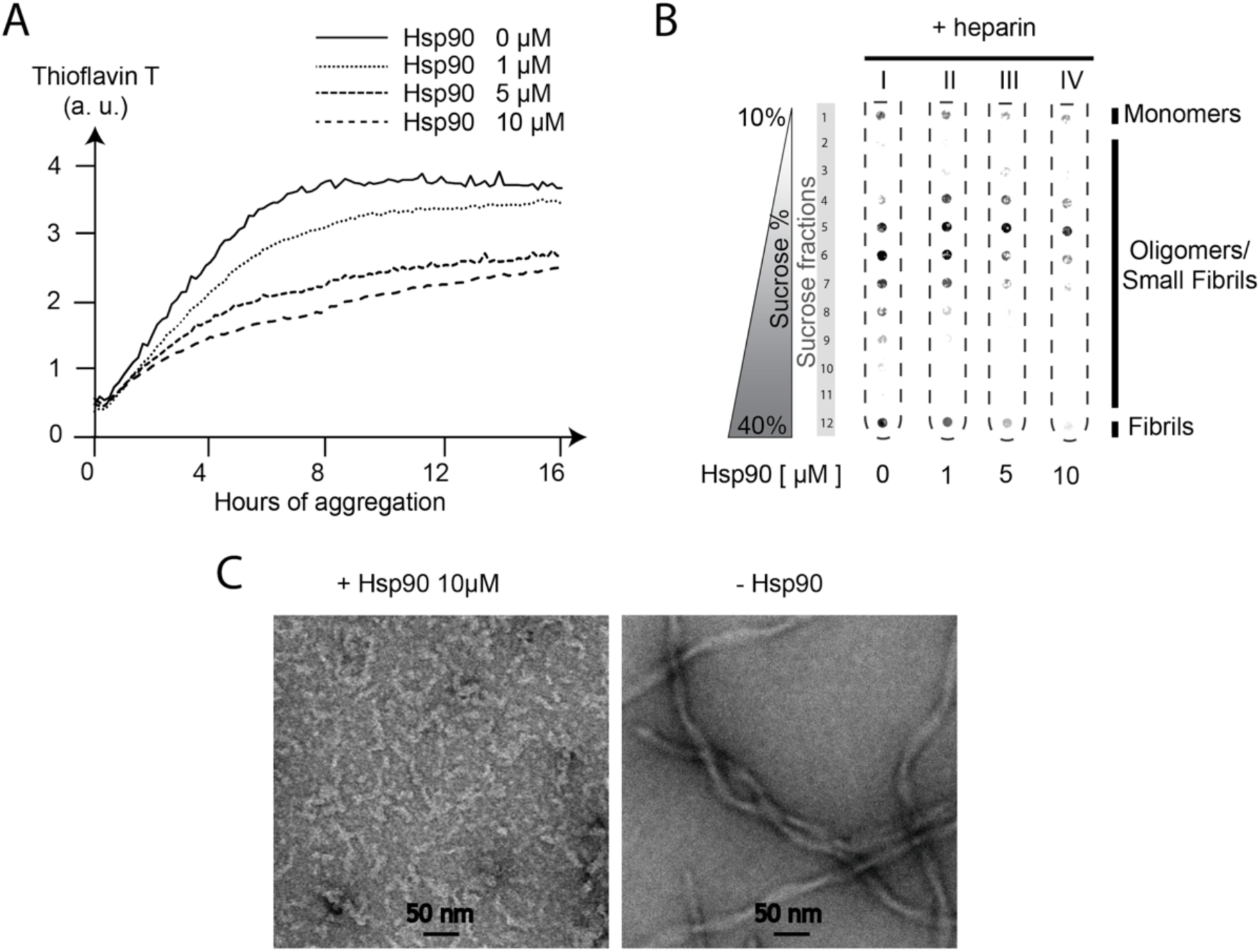
Hsp90 derails Tau aggregation toward non-fibrillar nano-aggregates. (A) Thioflavin T assay to detect decrease of amyloid content in the presence of increasing concentrations of Hsp90 (0, 1, 5, 10 /μM). Fluorescence is expressed as arbitrary unit (a. u.). (B) Density gradients to detect inhibition of fibril formation in the presence of increasing concentrations of Hsp90 (0, 1, 5, 10 /μM). Samples from **Fig. A** at 16 h were loaded directly onto density gradients and resolved by centrifugation for 2.5 h at 200,000 g. (C) Transmission Electron Microscopy of Tau-RD* aggregating samples in the presence or absence of 10 μM Hsp90 after 2 days of aggregation.

### Hsp90 modulates fibril interactome

Next, we wondered whether aggregates with different structural features should attract different binding partners. To address this question, we monitored Tau-RD* aggregation in the presence or absence of Hsp90 and described the changes in abundance of pulled-down interactors in these two conditions (**Fig. 6A**). Stunningly, none of the fibril-specific interactors were enriched, whereas many of them decreased in abundance for all functional clusters when Hsp90 modulated the aggregation (**Fig. 6B**). The only interactor that showed a significant enrichment was α-synuclein (SNCA), a well-known Hsp90 partner, suggesting that Hsp90 may co-aggregate with nano-aggregates. A subgroup of interactors, associated to Tau monomer but not to Tau fibrils (FBXO3, CTBP1, CTBP2 and POILIDIP2, **Fig. S1**) appeared in the presence of Hsp90. Also, the Hsp90-stalled nano-aggregates did not associate with COPI, in contrast to the ones formed in its absence. Taken together, these results suggest that Hsp90 drastically reshapes Tau aggregation-specific interactome, blocking the interactions associated to pathological Tau fibrils and moulding the interactome toward a more physiological state.

**Figure 6.**
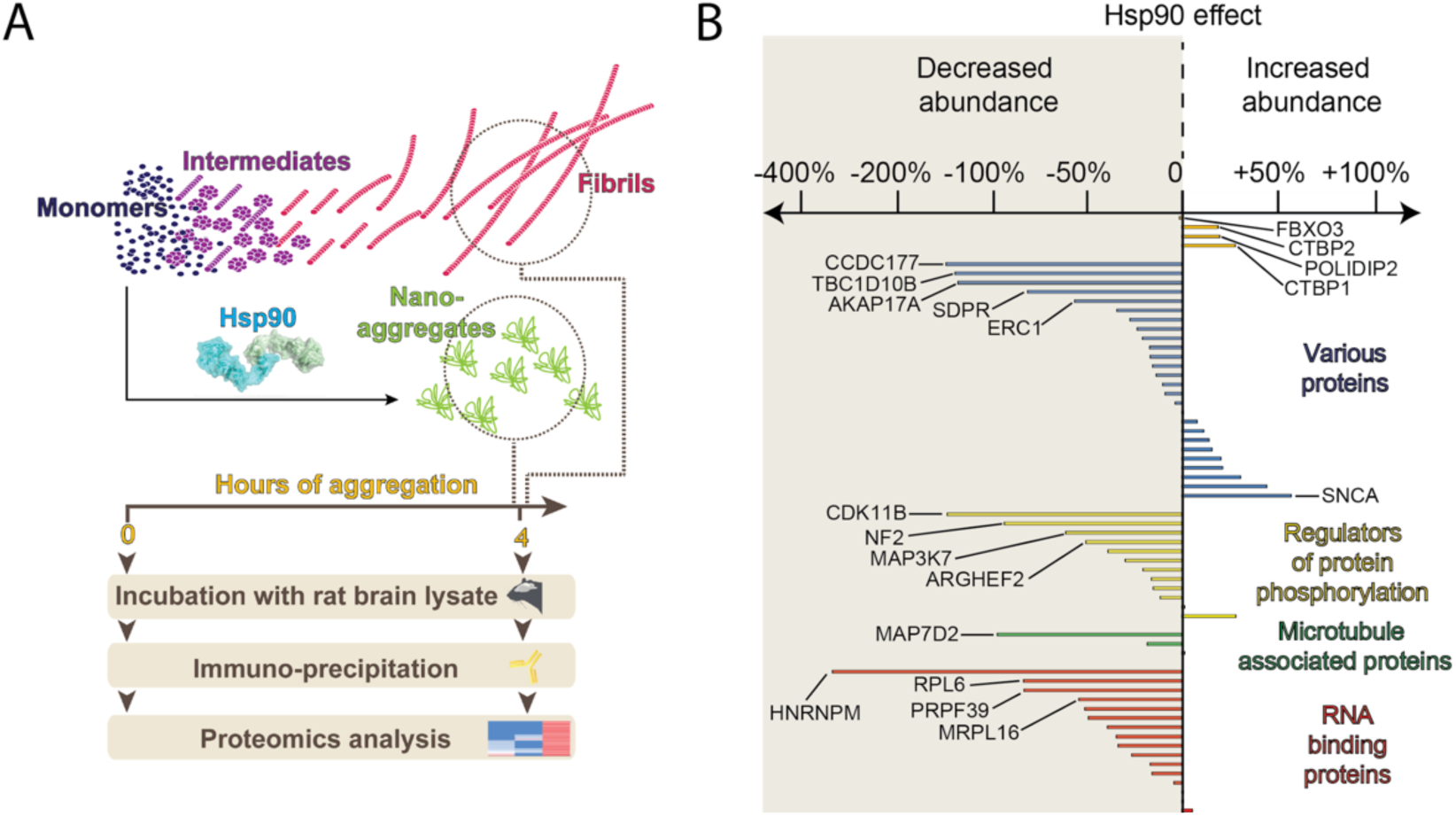
Hsp90 remodels Tau aggregation-specific interactome. (A) Experimental setup to compare interactome changes in the presence or absence of Hsp90. (B) Bar graph showing Hsp90-dependent modulation of Tau-RD* aggregation reflected as differences enrichments for specific proteins sequestered by Tau-RD* fibrils in the presence/absence of Hsp90.

## DISCUSSION

We tracked interactome changes associated to the formation of Tau fibrils, a hallmark of Alzheimer’s disease. Tau fibrils results from the aggregation of monomeric Tau, known to stabilize microtubules in neurons. As they form, Tau fibrils attract specific classes of abnormal interactors, mostly involved in RNA biology, regulation of protein phosphorylation and microtubule dynamics. Strikingly, these interactors share similar structural features, endowed with long disordered stretches with specific amino acidic bias. Arginine bias plays a crucial role, as Arginine-to-Lysine substitutions partially impairs the binding to Tau fibrils.

### Tau fibrils target RNA, cytoskeletal and phosphorylation dynamics

Our experimental setup focuses on the soluble fraction of the cytoplasm, the environment where fibrils naturally accumulate during the progression of Alzheimer (Wang and Mandelkow, 2016). Proteomics studies concerning Tau fibrils reveal interactomes of insoluble membrane-enriched fractions co-pelleted with Tau fibrils and their ER-associated components (Donovan et al., 2012; Meier et al., 2015). Here we provide data sets revealing interactions of Tau fibrils with the soluble components of the brain, to which the protein is exposed when aggregating in disease. Our setup accelerates the process of Tau aggregation from years *in vivo* to hours *in vitro* and highlights proteomic re-arrangements over time, adding a time dimension to proteomics studies. Thus, our study mimics the temporal dynamics of Tau aggregation spanning several decades from initial seeds to mature fibrils, with different aggregation stages having different reactivity with the cellular environment (**Fig. 2B**).

We fractionated Tau aggregates over time to overcome the problem that the dynamic nature of the aggregation process would preclude us from obtaining a defined, mono-disperse Tau nano-aggregate (Wang and Mandelkow, 2016). We show that early-stage aggregates specifically attract COPI components. These interactors decrease in abundance as fibrils form, highlighting the exchange of interactors as aggregation proceeds. Our data can explain Golgi fragmentation observed in neurons of Alzheimer brains without tangles (Stieber et al., 1996).

Our interactome data suggest that Tau fibrils may hijack three cellular processes: 1) RNA-related processes, 2) cytoskeletal dynamics via Microtubule Associated Proteins (MAPs) and 3) phosphorylation equilibria (**Fig. 2C**). Regarding RNA-related processes, the interactome of monomeric Tau is already highly enriched in components of the ribonucleoproteome (Gunawardana et al., 2015), and Tau fibrils have been shown to impair RNA translation (Meier et al., 2016). Our data suggest a mechanism where Tau fibrils attract additional proteins related to RNA processing, altering the balance between Tau and the ribonucleoproteome and thus impairing translation.

Tau fibrils target also MAPs, linked to cytoskeletal dynamics. MAPs have been shown to act together to regulate microtubule dynamics and cargo transport, with Map7-Tau imbalances disrupting axonal transport and neuronal growth (Monroy et al., 2018). Such imbalances in axonal transport are a common marker of Tau-related neurodegenerative disorders (Kneynsberg et al., 2017). Our data suggest a novel mechanism to explain such disorders, where Tau aggregation acts on two levels, first by promoting imbalances in the levels of different MAPs and second by actively sequestering other MAPs components, further exacerbating such imbalances.

Lastly, our data show how Tau fibrils attract a wide range of kinases and other protein phosphorylation regulators. Tau is hyperphosphorylated in Alzheimer, however the contribution of its phosphorylation to aggregation is yet to be understood (Goedert et al., 2017; Wang and Mandelkow, 2016). New structural insight proved that phosphorylated residues do not contribute significantly to assembly of mature fibrils (Fitzpatrick et al., 2017). Our data suggest that the effect of Tau fibrils on phosphorylation dynamics may be subtler and not related to their hyperphosphorylation, once again disturbing levels and activity of a range of phospho-regulators as in the case of RNA binding proteins and MAPs.

To summarize, scattered experimental evidences support the targeting of RNA, microtubule and phosphorylation dynamics by Tau fibrils. Our work shows how these processes are coherently linked by interactome re-wiring of Tau from its monomeric, physiological protomer to its polymeric, toxic aggregate.

### Analogies between mode-of-binding of Tau fibrils and proteins involved in phase separation

The structural analysis of Tau fibril-specific interactome revealed an abundance of disordered stretches compared to average disorder propensity of the human proteome. Intriguingly, the abundance of disorder stretches is a characteristic of interactors binding to Huntingtin fibrils (Ripaud et al., 2014) or to artificial proteins with propensity to form amyloid fibrils (Olzscha et al., 2011), suggesting that the preference for highly disordered binders is a general feature of amyloids.

We found a striking enrichment of Arginine and Methionine residues in disordered regions of Tau fibril binders, while Prolines and the hydrophobic residues Isoleucine, Valine and Phenylaniline were significantly depleted, highlighting a unique footprint of Tau fibrils binders. Lysine does not contribute to binding, despite their positive charge shared with Arginines, pointing to the role of Argine as π-stacker (Vernon et al., 2018). Of note, also the other strongly enriched residue in Tau fibril interactors, Methionine, can establish π-π interactions with aromatic rings (Valley et al., 2012). However, Arginines increased abundance suggests a predominant role of this residues in driving the binding to Tau fibrils. Thus, our work describes interactome re-arrangements not only in a qualitative manner but describe a key molecular component driving such re-arrangements, namely side chains of Arginines capable of engaging in π-π interactions.

π-stacking in disordered regions is also the fundamental force driving Liquid Liquid Phase Separation, a process that forms membrane-less compartmentalization within a cell (Alberti, 2017; Qamar et al., 2018). The number of Arginines is a key factor for phase separation propensity (Wang et al., 2018). So far, the role of n-stacking was showed only for physiologically relevant proteins. We turned the table, using non-physiological aggregates to show that their interactions are based on the same chemical principles driving phase separation. Our paradigm explains recent phenomenological data, showing co-influence of aggregation dynamics between Tau and phase-separating proteins (Maziuk et al., 2018; Vanderweyde et al., 2016). Strikingly, Tau fibril-specific interactome is enriched in proteins from the Heteronuclear Ribonucleoparticle family (**Fig. 2A** and **Fig. S2**), whose protein architecture promotes phase separation (Harrison and Shorter, 2017). Thus, Tau fibrils can attract phase-separating proteins, potentially increasing their local concentration and promoting their co-aggregation. Taken together, our data mechanistically support a tight cross-talk between physiological and pathological aggregates. In terms of fundamental chemical contacts, they are two sides of the same coin.

Interestingly, Arginine is enriched in several chaperone binding motifs, including Hsp70, J-domain co-chaperones, SecB and trigger factor (Knoblauch et al., 1999; Patzelt et al., 2001; Rüdiger et al., 1997; Rüdiger et al., 2001). We speculate that Arginine may have specific but not yet understood implications for protein homeostasis and aggregation. In this context it is interesting that the abundant cytosolic Hsp90 chaperone (Picard, 2002) may have an upstream effect in avoiding the engagement of Tau fibrils with their abnormal interactors by buffering these exposed stretches in the cytoplasm. Hsp90 may therefore act on two levels in the fight against Alzheimer-related Tau fibrils, first by actively blocking their formation and second by preventing fibrils binding to their abnormal interactors. Understanding how the manipulation of Hsp90 affect the biology of Tau fibrils may grant us a new weapon to fight Alzheimer’s Disease.

**Star * Methods**

**KEY RESOURCES TABLE**

| REAGENT OR RESOURCE | SOURCE | IDENTIFIER |
| --- | --- | --- |
| **ANTIBODIES** |  |  |
| ANTI-AMYLOID FIBRILS OC RABBIT POLYCLONAL ANTIBODY | Merck Millipore | AB2286 |
| ANTI-FLAG M2 MOUSE MONOCLONAL ANTIBODY | Sigma-Aldrich | F1804 |
| ANTI-HA 12CA5 MOUSE HYBRIDOMA | In-House | N/A |
| ANTI-FLAG® M2 MAGNETIC BEADS | Sigma-Aldrich | M8823 |
| ANTI-POLY HISTIDINE MOUSE MONOCLONAL ANTIBODY | Sigma-Aldrich | H1029 |
| DONKEY ANTI MOUSE IGG ALEXA FLUOR 680 | Invitrogen | A10038 |
| DONKEY ANTI MOUSE IGG IR DYE 800 CONJUGATED | Rockland | 610-732-002 |
| GOAT ANTI-RABBIT IR DYE 800NM CONJUGATED | Rockland | 611-132-002 |
| **BACTERIAL AND VIRUS STRAINS** |  |  |
| BL21 ROSETTA 2 CELLS | Novagen | 71402 |
| PET24A | Novagen | 69749 |
| PET23A | Novagen | 69745 |
| **CHEMICALS, PEPTIDES, AND RECOMBINANT PROTEINS** |  |  |
| 4X LAEMMLI SAMPLE BUFFER | Biorad | 161-0747 |
| ACETIC ACID 100% | Merck | 64-19-7 |
| ACETONITRILE | Merck | 100030 |
| ALBUMIN FROM CHICKEN EGG | Sigma-Aldrich | A7641 |
| AMMONIUM BICARBONATE | Sigma-Aldrich | 09830 |
| AMPICILLIN SODIUM SALT | Sigma-Aldrich | A9518 |
| CHLORAMPHENICOL | Sigma-Aldrich | C0378 |
| COMPLETE PROTEASE COCKTAIL INHIBITORS | Roche | CO-RO |
| COMPLETE PROTEASE EDTA FREE COCKTAIL INHIBITORS | Roche | COEDTAF-RO |
| DTT | Thermo Fischer Scientific | R0862 |
| GELCODE BLUE STAIN REAGENT | Thermo Fischer Scientific | 24590 |
| HEPARIN LOW MOLECULAR WEIGHT | Santa Cruz Biotech | 1-8-9041 |
| HEPES | Sigma-Aldrich | H3375 |
| HEPES SODIUM SALT | Sigma-Aldrich | H7006 |
| IMIDAZOLE | Sigma-Aldrich | 56749 |
| IPTG | Thermo Fischer Scientific | 15529019 |
| KANAMYCIN SULFATE FROM *STREPTOMYCES KANAMYCETICUS* | Sigma-Aldrich | K4000 |
| KCL | Carl Roth | 6781.2 |
| LI-COR ODYSSEY BLOCKING BUFFER (TBS) | Westburg | LI 927-50000 |
| IMIDAZOLE | Sigma-Aldrich | 56749 |
| IDOACETAMIDE | Merck | 144-48-9 |
| METHANOL | Merck | 106009 |
| NACL | Carl Roth | HN00.3 |
| NONIDET® P40 | AppliChem | A1694 |
| **NYCODENZ AG** | Axis-Shield | 1002424 |
| **SDS SOLUTION, 20 %** | Serva | 20767.03 |
| **SODIUM AZIDE** | Sigma-Aldrich | S8032 |
| **SODIUM FLUORIDE** | Sigma-Aldrich | S7920 |
| **SODIUM PHOSPHATE MONOBASIC** | Sigma-Aldrich | S3139 |
| **SUCROSE** | Sigma-Aldrich | S0389 |
| **THIOFLAVIN T** | Sigma-Aldrich | 2390-54-7 |
| **TRITON X-100** | Sigma-Aldrich | 9002-93-1 |
| **TRIZMA BASE** | Sigma-Aldrich | T6066 |
| **TRYPSIN** | Promega | V5111 |
| **ULP1** | In-house | 12588018 |
| **B-MERCAPTOETHANOL** | Sigma-Aldrich | M3148 |
| **CRITICAL COMMERCIAL ASSAYS** |  |  |
| **BCA ASSAY** | Thermo Fischer Scientific | 23250 |
| **WIZARD PLUS SV MINIPREPS** | Promega | A1460 |
| **RECOMBINANT DNA** |  |  |
| **PET24A HUMAN SUMO-TAU-RD\*** | In-house | NA |
| **PET24A RAT MAP7 SMT<sub>6</sub>-HA-M1-S227 WT** | In-house | NA |
| **PET24A RAT MAP7 SMT<sub>6</sub>-HA-M1-S227 WT** | In-house | NA |
| **PET23A HUMAN HISHSP90β** | In-house | NA |
| **SOFTWARE AND ALGORITHMS** |  |  |
| **CRAPOME** | www.crapome.org | NA |
| **IMAGE STUDIO 2.10.1** | LI-COR | NA |
| **PERSEUS** | https://www.nature.com/articles/nmeth.3901#plugin-architecture | NA |
| **PRISM7** | Graphpad | NA |
| **PROTEOME DISCOVERER 1.4** | ThermoScientific | NA |
| **R** | https://www.r-project.org/ | NA |
| **XCALIBUR™ SOFTWARE** | ThermoScientific | NA |
| **OTHER** |  |  |
| **4–12% CRITERION XT BIS-TRIS PROTEIN GEL** | Biorad | 3450123 |
| **96-WELL POLYPROPYLENE DEEP WELL STORAGE PLATES** | Thermo Fischer Scientific | 260251 |
| **96-WELL GREINER PLATES, POLYSTYRENE** | Sigma-Aldrich | M3061 |
| **AGILENT 1290 INFINITY LC** | Agilent Technologies |  |
| **CIRCULAR DICHROISM SPECTROPOLARIMETER J-810** | JASCO | 0302-0407A |
| **DYNAMAG -2 MAGNET** | Invitrogen | 10723874 |
| **HITRAP HEPARIN COLUMN HIGH PERFORMANCE** | GE Healthcare | 17-0407-01 |
| **POROS20HQ ANION EXCHANGE COLUMN** | Thermo Fischer Scientific | 1232906 |
| **POROS20HQ CATION EXCHANGE COLUMN** | Thermo Fischer Scientific | 1332906 |
| **POROS20MC AFFINITY PURIFICATION COLUMN** | Thermo Fischer Scientific | 1542906 |
| **POROSHELL 120 EC C18 COLUMNS** | Agilent Technologies | 699775-922 |
| **ND-1000 UV/VIS SPECTROPHOTOMETER** | NanoDrop Technologies | S137007 |
| **Q EXACTIVE ORBITRAP MASS SPECTROMETER** | Thermo Fischer Scientific | IQLAAEGAA<br>PFALGMAZR |
| **REPROSIL-PUR BASIC C18, 3 μM** | Dr.Maisch | r13.b9. |
| SPECTRA/POR DIALYSIS<br>MEMBRANE MWCO 12-14 KDA | SpectrumLaboratories | 132703 |
| SPECTRA/POR DIALYSIS<br>MEMBRANE MWCO 3.5 KDA | SpectrumLaboratories | 132724 |
| SPECTRAMAX I3 | Molecular Devices | NA |
| SORVALL WX ULTRA SERIES<br>CENTRIFUGE WX80 | Thermo Fischer Scientific | 75000080 |
| VIVASPIN 6 BUFFER EXCHANGE<br>FILTER | Sartorius | VS0612 |
| WHATMAN PROTRAN<br>NITROCELLULOSE<br>MEMBRANE | Sigma-Aldrich | Z670952 |

## CONTACT FOR REAGENT AND RESOURCE SHARING

Further information and requests for resources and reagents should be directed to and will be fulfilled by the Lead Contact, S. G. D. Rudiger.

## EXPERIMENTAL MODEL AND SUBJECT DETAILS

### In Vitro Studies

Human Tau-RD* and rat HA-Map7 truncations (M1-S227, both wild type and R_12_K), were cloned in pET24a vectors and expressed in *E. coli* (BL21 Rosetta2).

Human Hsp90β was cloned in pET23a vector and expressed in *E. coli* (BL21 Rosetta2).

### Animals

All experiments were approved by the DEC Dutch Animal Experiments Committee (Dier Experimenten Commissie), performed in line with institutional guidelines of Utrecht University and were conducted in agreement with Dutch law (Wet op de Dierproeven, 1996) and European regulations (Directive 2010/63/EU). Female pregnant Wister rats were obtained from Janvier Laboratories.

## METHOD DETAILS

### Cloning, expression and purification

#### Purification of Tau-RD*

We overproduced N-terminally FLAG-tagged (DYKDDDDK) Human Tau-RD (Q244-E372, also referred to as K18, with pro-aggregation mutation ΔK280) recombinantly in *E. coli* BL21 Rosetta 2 (Novagen), with additional removable N-terminal His6-Smt-tag (MGHHHHHHGSDSEVNQEAKPEVKPEVKPETHINLKVSDGSSEIFFKIKKTTPL RRLMEAFAKRQGKEMDSLRFLYDGIRIQADQTPEDLDMEDNDIIEAHREQIGGJ. Cells were harvested, flash-frozen in liquid nitrogen and stored at −80°C until further usage. Pellets were thawed in a water bath at 37° C and resuspended in 50 mM HEPES-KOH pH 8.5 (Sigma-Aldrich), 50 mM KCl (Sigma-Aldrich), ½ tablet/50 ml EDTA-free protease inhibitor (Roche), 5 mM β-mercaptoethanol (Sigma-Aldrich). Cells were disrupted by an EmulsiFlex-C5 cell disruptor (Avestin). Lysate was cleared by centrifugation, filtered with a 0.22 μm polypropylene filter (VWR) and supernatant was purified using an ÄKTA Purifier chromatography system (GE Healthcare). First, protein was loaded onto a POROS 20MC (Thermo Fischer Scientific) affinity purification column in 50 mM HEPES-KOH pH 8.5, 50 mM KCl, 5 mM β-mercaptoethanol, eluted with a 0-100% linear gradient (5 Column Volumes, CV) of 1 M imidazole. Fractions of interest were collected and concentrated in a buffer concentrator (Vivaspin, cut-off 10 kDa) to final volume of 3 ml. The concentrated sample was desalted with a PD-10 desalting column (GH Healthcare) in 50 mM HEPES-KOH pH 8.5, ½ tablet/50 ml Complete protease inhibitor (Roche)) and 5 mM p-mercaptoethanol. The His6-Smt tag was removed by Ulp1 treatment, shaking at 4°C over night. Next day, protein was loaded onto a POROS 20HS (Thermo Fischer Scientific) cation exchange column equilibrated with 50 mM HEPES-KOH pH 8.5. Protein was eluted with a 0-100% linear gradient (15 CV) of 2 M KCl (Carl Roth). Fractions of interest were collected and loaded onto a HiLoad 26/60 Superdex 200 pg (GE Healthcare Life Sciences) size exclusion column equilibrated with aggregation buffer (25 mM HEPES-KOH pH 7.5, Complete Protease Inhibitors (1/2 tablet/50 ml), 75 mM KCl, 75 mM NaCl and 10 mM DTT). Fractions of interest were further concentrated to the desired final concentration using a concentrator (Vivaspin, cut-off 5 kDa). Protein concentration was measured using an ND-1000 UV/Vis spectrophotometer (Nanodrop Technologies) and purity was assessed with SDS-PAGE. Protein was aliquoted and stored at −80°C.

#### Purification of MAP7 truncations

We overproduced N-terminally HA-tagged (YPYDVPDYA) rat map 7 truncations (M1-S227, both wild type and R_12_K), recombinantly in *E. coli* BL21 Rosetta 2 (Novagen), with additional removable N-terminal His6-Smt-tag. Cells were harvested, flash-frozen in liquid nitrogen and stored at −80°C until further usage. Pellets were thawed in a water bath at 37°C and resuspended in 50 mM phosphate buffer pH 8 (Sigma-Aldrich), 150 mM KCl (Sigma-Aldrich), ½ tablet/50 ml EDTA-free protease inhibitor (Roche), 5 mM β-mercaptoethanol (Sigma-Aldrich). Cells were disrupted by an EmulsiFlex-C5 cell disruptor (Avestin). Lysate was cleared by centrifugation, filtered with a 0.22 ^m polypropylene filter (VWR) and supernatant was purified using an ÄKTA Purifier chromatography system (GE Healthcare). First, protein was loaded onto a column with Ni-IDA resin. Proteins were eluted by 250 mM imidazole (Sigma-aldrich) in phosphate buffer (150 mM, NaCl (Sigma-Aldrich), pH=8.0) in presence of Complete Protease Inhibitors (1 tablets/100 ml) (Roche) and 5 mM βmercaptoethanol (Sigma-Aldrich).The His_6_-Smt tag was removed by Ulp1 treatment at 4°C over night, while dialyzed against 50mM phosphate buffer (pH=8.0) and 5 mM βmercaptoethanol with a 6kDa cut-off membrane (SpectrumLaboratories). Next day, protein was loaded onto a POROS 20HS (Thermo Fischer Scientific) cation exchange column equilibrated with 50mM phosphate buffer (pH=8.0) and 5 mM βmercaptoethanol. Protein was eluted with a 0-100% linear gradient (15 CV) of 2 M KCl (Carl Roth). Fractions of interest were further buffer exchanged againt 25 mM HEPES buffer (pH=7.5, salt: 75 mM KCl; 75 mM NaCl) Vivaspin, cut-off 6 kDa and further concentrated using a concentrator (Vivaspin, cut-off 5 kDa). Protein concentration was measured using an ND-1000 UV/Vis spectrophotometer (Nanodrop Technologies) and purity was assessed with SDS-PAGE. Protein was aliquoted and stored at −80°C.

#### Protein purification of Hsp90

Hsp90 was purified as previously described (Radli et al., 2017). We overproduced N-terminally His6-tagged human Hsp90β recombinantly in *E. coli* BL21 Rosetta 2 (Novagen). Cells were harvested, flash-frozen in liquid nitrogen and stored at −80°C until further usage. Pellets were thawed in a water bath at 37°C and resuspended in 12.5 mM sodium phosphate buffer pH 6.8 (Sigma-Aldrich), 75 mM KCl (Sigma-Aldrich), ½ tablet/50 ml EDTA-free protease inhibitor (Roche), 5 mM β-mercaptoethanol (Sigma-Aldrich). Cells were disrupted by an EmulsiFlex-C5 cell disruptor (Avestin). Lysate was cleared by centrifugation, filtered with a 0.22 μm polypropylene filter (VWR) and supernatant was purified using an ÄKTA Purifier chromatography system (GE Healthcare). First, protein was loaded onto a POROS 20MC (Thermo Fischer Scientific) affinity purification column in 50 mM sodium phosphate buffer pH 8.0, 400 mM KCl, 5 mM β-mercaptoethanol eluted with a 0-100% linear gradient (5 Column Volumes, CV) of 1 M imidazole. Peak was loaded onto a POROS 20HQ (Thermo Fischer Scientific) anion exchange column equilibrated with 25 mM sodium phosphate buffer pH 7.2 and 5 mM β-mercaptoethanol. Protein was eluted with a 0-100% linear gradient (15 CV) of 2 M KCl. Fractions of interest were then loaded onto a HiTrap heparin column High Performance (GE Healthcare) column equilibrated with 25 mM sodium phosphate buffer pH 7.2 and 5 mM β-mercaptoethanol. Protein was eluted with a 0-100% linear gradient (15 CV) of 2 M KCl and peak was buffer exchanged against 25 mM sodium phosphate buffer pH 7.2, 150 mM KCl, 150 mM NaCl and 5 mM βmercaptoethanol. Protein concentration was measured using an ND-1000 UV/Vis spectrophotometer (Nanodrop Technologies) and purity was assessed with SDS-PAGE. Protein was aliquoted and stored at −80°C.

### Formation of Tau fibrils

Monomeric Tau-RD* aggregated in aggregation buffer and Heparin Low Molecular Weight (Santa Cruz Biotech), concentrations depending on the experiment and Tau:heparin ratio always kept 4:1. Aggregation was performed at 37°C, shaking at 180 rpm, and aliquots were flash frozen at timepoints indicated in the text. Aliquots were then thawed on ice for downstream applications.

### Preparation of density gradients

Density gradients were prepared according to an established procedure (Stone, 1974). Gradients were formed by dissolving 10% and 40% sucrose (Sigma-Aldrich), in 25 mM HEPES pH 7.5, 75 mM KCl, 75 mM NaCl. Gradients were set up in polyallomer centrifuge tubes (Beckmann) by filling them to half hight with 40% sucrose and topping them up with an equal amount of 10% sucrose. Gradients were formed by tilting the tubes horizontally for 3 h at room temperate and then tilting them back to vertical position. Tubes were stored overnight at 4°C and samples were loaded as described for each experiment.

### Sample preparation for density gradients

**Fig. 1A.** 2.5 μl of Tau-RD* aggregates (37 μM of monomers) at different aggregation stages (time of aggregations: 0, 1, 8, 24 h) were loaded onto density tubes and subjected to centrifugation, 2.5 h at 200,000 g.

**Fig. 4D**. 2.5 μl of Tau-RD* fibrils (20 μM of monomers, timepoint 24 hours) reacted for 1 h at 37°C with either 7.5 μl of aggregation buffer or 7.5 μl of aggregation buffer plus 0.5 μM rat HA-Map7 M1-S227, either wt or R_12_K. Samples were then entirely loaded loaded onto density tubes and subjected to centrifugation, 2.5 h at 200000g.

**Fig. 5B**. 10 μl of Tau-RD* (20 μM of monomers), aggregated for 16 hours in the plate reader Spectramax i3 (Molecular Devices) in the presence of increasing concentration of Hsp90 (0, 1, 5 and 10 μM) in aggregation buffer, were loaded completely onto density tubes and subjected to centrifugation, 2.5 h at 200000g.

### Dot blot analysis of density gradients

Dot blot was performed using a dot blot apparatus (BioRad) and nitrocellulose membrane 0.1 μM (Sigma-Aldrich) washed with PBS. Twelve fractions were manually collected for each tube into a 96-deep well (Thermo Fisher Scientific), and each dot-blot well was filled with 150 μl of fraction using a multi-pipette. which was pulled through by applying vacuum after 10 minutes of incubation with the membrane at room temperature. Nitrocellulose membranes were blocked with PBS blocking buffer (LI-COR) and incubated with primary antibody, either monoclonal anti-FLAG M2 (F3165, Sigma Aldrich, working dilution 1:1000) or anti-HA 12CA5 mouse hybridoma (produced in-house), at room temperature for 1 h. After three washes with PBS, secondary antibody Donkey anti Mouse IgG IR Dye 800 conjugated (610-732-002, Rockland, 1:5000) was added at room temperature for 45 minutes. After additional 2 washes with PBS-T and one final wash with PBS, detection was performed using Odyssey CLx (LI-COR). Imaging was performed via Image Studio Lite (LI-COR).

### Transmission Electron Microscopy

Specimens were prepared for Transmission Electron Microscopy using a negative staining procedure. Briefly, a 5 μl drop of sample solution was adsorbed to a glow-discharged (twice for 20 s, on a Kensington carbon coater) pioloform-coated copper grid, washed five times on drops of deionized water, and stained with two drops of freshly prepared 2.0% Uranyl Acetate, for 1 and 5 minutes, respectively, and subsequently air dried. Samples were imaged at room temperature using a Tecnai T20 LaB6 electron microscope operated at an acceleration voltage of 200 kV and equipped with a slow-scan Gatan 4K x 4K CCD camera. Images were acquired at a defocus value of 1.5 μm. Magnification/pixel size on specimen level: **Fig. 1A** 62.000 times/0.178 nm/pix; **Fig. 5C** right panel 62.000 times, 0.178 nm/pix; **Fig. 5C** right panel 100.000 times, 0.110 nm/pix

### Rat brain extracts preparation

Rat brain extracts were obtained from female adult rats and homogenized in 10x volume/weight in tissue lysis buffer (50 mM Tris HCl, 150 mM NaCl, 0.1% SDS, 0.2% NP-40, and protease inhibitors). Brain lysates were centrifuged at 16.000 g for 15 min at 4°C, and the supernatant was used for the affinity purification-mass spectrometry experiments.

### Affinity Purification-Mass spectrometry (AP-MS) using anti-FLAG beads on brain extracts

Tau-RD* monomers, oligomers or fibrils (37 μM monomeric concentration, with or without 37 μM Hsp90, depending on the experiment) were incubated for 1h at 4°C with FLAG beads (Sigma) previously blocked in chicken egg albumin (Sigma). Beads were then separated using a magnet (Dynal, Invitrogen) and washed three times with aggregation buffer to remove unbound Tau-RD* and excess of albumin. Beads conjugated with the Tau-RD* aggregated proteins were then incubated with brain extracts for 1h at 4°C. Beads were then washed in washing buffer (20 mM Tris HCl, 150 mM KCl, 0.1% TritonX-100) for five times to remove aspecific neuronal proteins. For MS analysis, beads were then resuspended in 15 μl of 4x Laemmli Sample buffer (Biorad), boiled at 99°C for 10 minutes and supernatants were loaded on 4-12% Criterion XT Bis-Tris precast gel (Biorad). The gel was fixed with 40% methanol and 10% acetic acid and then stained for 1h using colloidal coomassie dye G-250 (Gel Code Blue Stain, Thermo Scientific). Briefly, each lane from the gel was cut into three pieces and placed in 1.5-ml tubes. They were washed with 250 μl of water, followed by 15-min dehydration in acetonitrile. Proteins were reduced (10 mM DTT, 1 h at 56°C), dehydrated and alkylated (55 mM iodoacetamide, 1 h in the dark). After two rounds of dehydration, trypsin was added to the samples (20 μl of 0.1 g/l trypsin in 50 mM Ammoniumbicarbonate) and incubated overnight at 37°C. Peptides were extracted with acetonitrile, dried down and reconstituted in 10% formic acid prior to MS analysis.

### Mass spectrometry analysis

All samples were analyzed on an Orbitrap Q-Exactive mass spectrometer (Thermo Fisher Scientific) coupled to an Agilent 1290 Infinity LC (Agilent Technologies). Peptides were loaded onto a trap column (Reprosil pur C18, Dr. Maisch, 100 μm x 2 cm, 3 μm; constructed in-house) with solvent A (0.1% formic acid in water) at a maximum pressure of 800 bar and chromatographically separated over the analytical column (Poroshell 120 EC C18, Agilent Technologies, 100 μm x 50 cm, 2.7 μm) using 90 min linear gradient from 7-30% solvent B (0.1% formic acid in acetonitrile) at a flow rate of 150 nl/min. The mass spectrometers were used in a data-dependent mode, which automatically switched between MS and MS/MS. After a survey scan from 350-1500 m/z the 10 or 20 most abundant peptides were subjected to HCD fragmentation. MS spectra were acquired in high-resolution mode (R > 30,000), whereas MS2 was in high-sensitivity mode (R > 15,000).

### Gene Ontology analysis

Proteins were classified using the enrichment analysis tool provided by Gene Ontology Consortium (GO Consortium, 2017 (Ashburner et al., 2000)).

### Circular Dichroism

HA-tagged (YPYDVPDYA) rat map 7 truncations (M1-S227, both wild type and R_12_K) were buffer exchanged against chloride-free phosphate buffer pH 7.5, 150 mM NaF (Sigma-Aldrich), to an estimated concentration of 0.05 mg/ml. Spectra were obtained on a Circular Dichroism Detector (JASCO), range from 180 nm to 250 nm, cut off at high tension over 700 V.

### ThioflavinT aggregation assay

Aggregation of Tau-RD* (20 μM stock) in aggregation buffer (total volume 100 μl) was stimulated by 5 μM heparin Low Molecular Weight in the presence of 60 μM ThioflavinT (Sigma) in transparent, lidded Greiner 96-well plates (Sigma-Aldrich). To assess the impact of Hsp90, samples were supplied with 0, 1, 5 or 10 μM of Hsp90. Fluorescent spectra were recorded every 10 minutes for 16 hours with a SpectraMax i3 (Molecular Devices).

## QUANTIFICATION AND STATISTICAL ANALYSIS

### Disorder prediction and Amino acidic bias

Disordered residues in Tau interactome protein were predicted using MetaDisorder predictor (Kozlowski and Bujnicki, 2012). We defined disorder percentage of each interactor as the amount of disordered amino acids divided by total amount of amino acids.

To calculate aminoacidic biases in disordered regions, disordered regions of each aggregation-specific interactor were considered as a single string of amino acids. The frequency of each amino acid in each disordered string was then calculated and compared to its frequency in the disordered regions of the whole proteome (Pentony and Jones, 2010). The ratio of calculated over predicted frequency was then computed and expressed as a log2 function for the whole set of aggregation-specific interactors, sorted per amino acid, with residues ordered by their abundance in the human proteome.

### Boxplot and graphs of Fig. 3 were created with SPSS

Statistical significance of **Fig. 3B** and **Fig. 3C** was tested using the Wilcoxon paired non-parmatric t-test on measured paired to predicted data for each protein.

### Mass spectrometry quantification

To infer protein abundancy of each individual protein co-purified with Tau-RD*, we relied on total numbers of PSMs; PSMs of each identified protein were then normalized on the PSMs of the purified Tau-RD* in each condition. A ratio was then calculated between normalized PSMs of the protein co-purified with Tau-RD* at different time point of aggregation and normalized PSMs of the same protein at 0 h without heparin. Ratios were then transformed into log2. Crapome (Mellacheruvu et al., 2013) was used to analyze Tau-RD* interacting binding proteins in three biological replicas. Proteins co-precipitated with Tau-RD* at 4 h were compared with proteins co-precipitated with Tau-RD* at 0 h without heparin. Proteins with a Fold Change calculation > 2 were considered significant proteins sequestered by Tau-RD* aggregates. This unbiased analysis for scoring AP-MS data generated a selection of proteins characterized by higher binding affinity for fibrillar species (**Fig. 2D**, **Fig. S2**). Same selected proteins are also shown in **Fig. 6B** to highlight how Tau-RD* interactomes change upon incubation with Hsp90. Proteins co-precipitated with Tau-RD* at 0h were compared with controls (FLAG-beads). Proteins with a Fold Change calculation > 2 were considered significant proteins sequestered by Tau-RD* monomers. This unbiased analysis for scoring AP-MS data, generated a selection of proteins characterized by higher binding affinity for monomeric species (**Fig. 2C**, **Fig. S1**). Data analysis was conducted using Perseus or R, hierarchical clustering was performed within Perseus using Euclidian distance. **Fig. 2B**: Quantifications of proteins detected at t = 0, 1, 8 and 24 h in replica 3 are represented in the heat map. **Fig. 2C**: Heat map shows an unbiased selection of proteins co-precipitating with Tau-RD* monomers without heparin (t_0-_) compared to control pull downs (empty FLAG-beads). Only proteins with a Fold Change calculation > 3 (FC, Crapome; by averaging the spectral counts across the selected controls) were considered enriched in Tau-RD* monomers without heparin (t_0-_) compared to controls in the 3 different biological replicas. Values shown in the heat map refer to time points t_0_, t_1_, t_8_ and t_24_ of replica 3. Fig. S1: Interactors of Tau-RD* monomers selected in **Fig. 2C** are shown with their relative quantifications across all time points in 3 biological replicas. **Fig. 2D** Heat map shows an unbiased selection of proteins specifically co-precipitating with Tau-RD* aggregates (t_4_) compared to Tau-RD* monomers without heparin (t_0-_). Only proteins with a Fold Change calculation > 2 (FC, Crapome; by averaging the spectral counts across the selected controls) were considered enriched in Tau-RD* monomers without heparin (t_0-_) compared to controls in the 3 different biological replicas. Values shown in the heat map refer to time points t_0_, t_1_, t_8_ and t_24_ of replica 3. **Fig. S2:** Full list of interactors of Tau-RD* oligomers and fibrils (t_4_) shown in **Fig. 2D** is shown with their relative quantifications across all time points in 3 biological replicas. **Fig. 6B**: Bar graph shows same selected proteins of Fig. S2 (significant proteins interacting with Tau-RD* aggregates, t_4_) and their average quantifications in Tau-RD* (t_4_) compared to Tau-RD* + Hsp90 (t_4_) across 3 different biological replicas. Ratios of normalized PSMs of interactors in Tau-RD* - Hsp90 / normalized PSMs of interactors in Tau-RD*+Hsp90 are graphically represented and shown as percentages. Negative values indicate a decreased binding affinity in presence of Hsp90 while positive ones indicate a higher affinity. Quantification of selected interactors of Tau-RD* monomers without heparin t_0-_ (Fbxo3, Poldip2, Ctbp1, Ctbp2) is also included as additional control. Most of the interactors of Tau-RD* aggregates reduce their binding affinity in presence of Hsp90 whereas Fbxo3, Poldip2, Ctbp1 and Ctbp2 do not.

## DATA AND SOFTWARE AVAILABILITY

### Software

#### Mass spectrometry data analysis

For data analysis, the raw data files were converted to *.mgf files using Proteome Discoverer 1.4 software (Thermo Fisher Scientific). Database search was performed using the Uniprot rat database and Mascot (version 2.5.1, Matrix Science, UK) as the search engine. Carbamidomethylation of cysteines was set as a fixed modification and oxidation of methionine was set as a variable modification. Trypsin was set as cleavage specificity, allowing a maximum of 2 missed cleavages. Data filtering was performed using a percolator (Kall et al., 2007), resulting in 1% false discovery rate (FDR). Additional filters were search engine rank 1 and mascot ion score >20.

## SUPPLEMENTARY FIGURES

**Figure S1.**
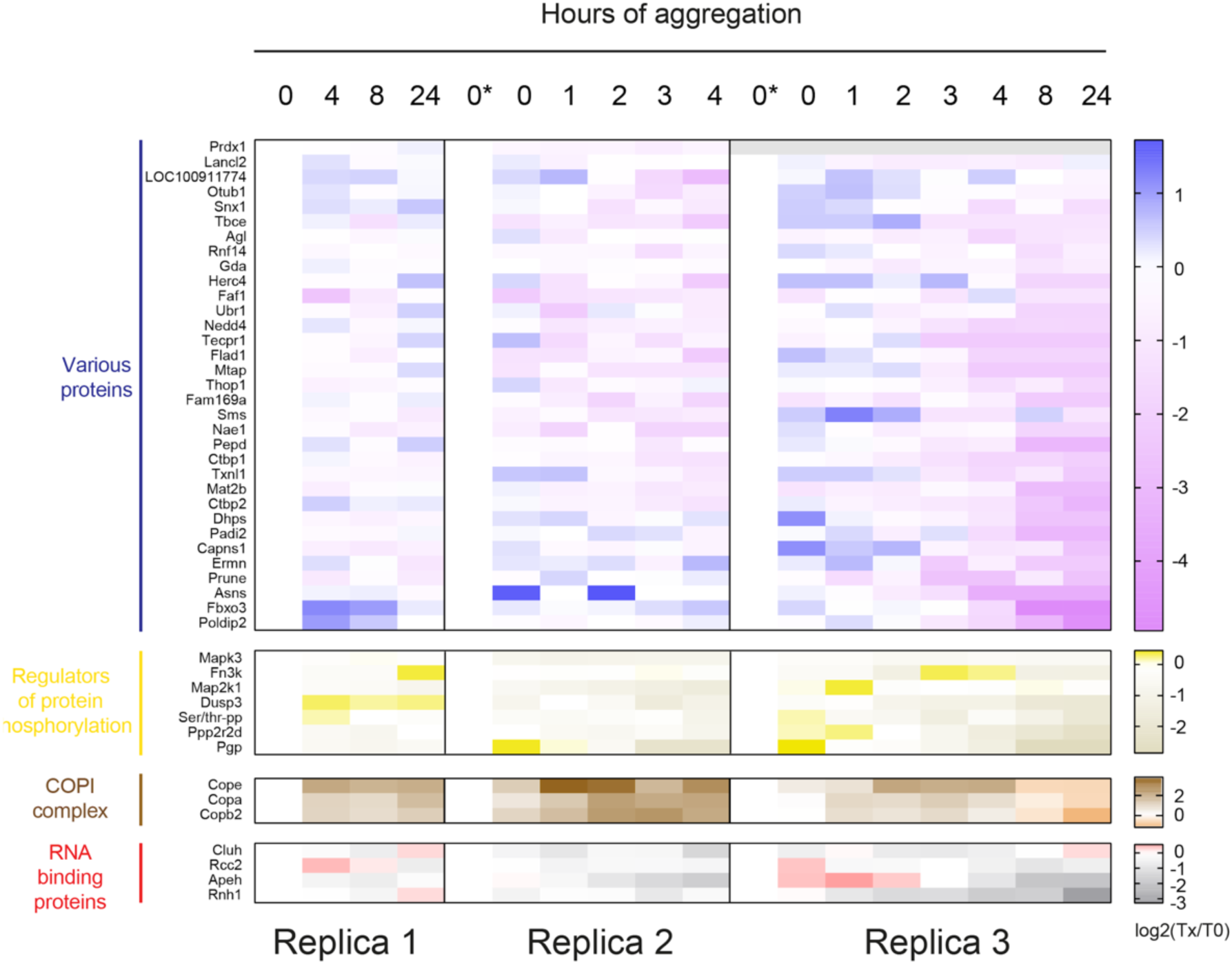
Interactome rewiring of monomeric-specific interactors, 3 independent biological replicas. Heat map showing an unbiased selection of Tau-RD* lost proteins upon aggregation. Only proteins with a Fold Change calculation > 2 (FC-A, Crapome; by averaging the spectral counts across the selected controls) were considered enriched in Tau-RD* monomers compared to control empty beads in the 3 different replicas.

**Figure S2.**
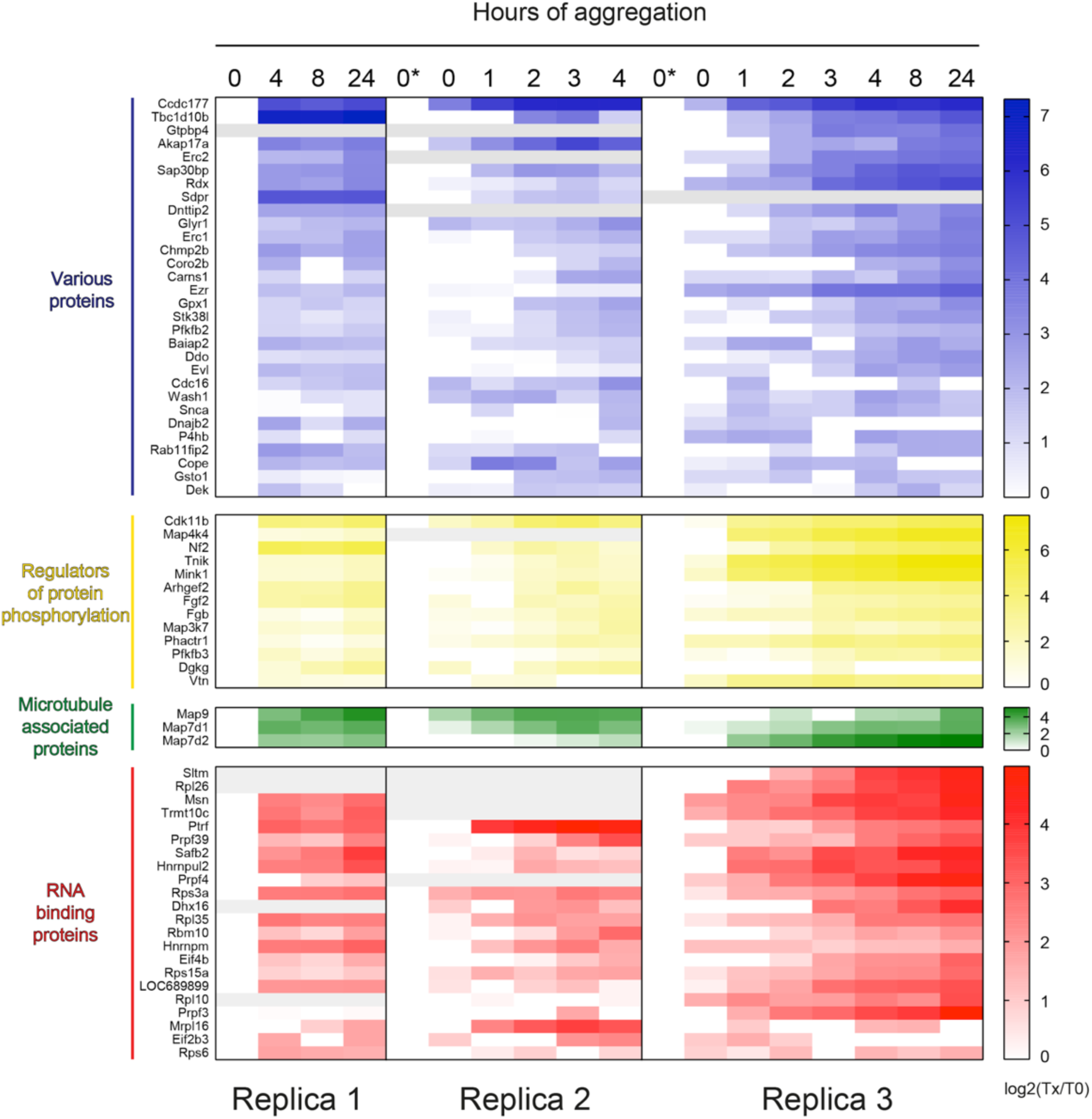
Interactome rewiring of fibrillar-specific interactors, 3 independent biological replicas. Heat map showing an unbiased selection of Tau-RD* sequestered proteins upon aggregation. Only proteins with a Fold Change calculation > 2 (FC-A, Crapome; by averaging the spectral counts across the selected controls) were considered enriched in Tau-RD* aggregates compared to Tau-RD* monomers in the 3 different replicas.

